# Development of an optimized and scalable method for isolation of umbilical cord blood-derived small extracellular vesicles for future clinical use

**DOI:** 10.1101/2020.12.18.423416

**Authors:** R.M.S. Cardoso, S.C. Rodrigues, C. F. Gomes, F.V. Duarte, M. Romao, E.C. Leal, P.C. Freire, R. Neves, J. Simões-Correia

## Abstract

Extracellular vesicles (EV) are a promising therapeutic tool in regenerative medicine. These particles were shown to accelerate wound healing, through delivery of regenerative mediators, such as microRNAs. Herein we describe an optimized and up-scalable process for the isolation of EV smaller than 200 nm (sEV), secreted by umbilical cord blood mononuclear cells (UCB-MNC) under ischemic conditions and propose quality control thresholds for the isolated vesicles, based on the thorough characterization of their protein, lipid and RNA content.

Ultrafiltration and size exclusion chromatography (UF/SEC) optimized methodology proved superior to traditional ultracentrifugation (UC), regarding production time, standardization, scalability, and vesicle yield. Using UF/SEC, we were able to recover approximately 400 times more sEV per mL of media than with UC, and up-scaling this process further increases EV yield by about 3-fold. UF/SEC-isolated sEV display many of the sEV/exosomes classical markers and are enriched in molecules with anti-inflammatory and regenerative capacity, such as hemopexin and miR-150. Accordingly, treatment with sEV promotes angiogenesis and extracellular matrix remodeling, *in vitro. In vivo*, UCB-MNC-sEV significantly accelerate skin regeneration in a mouse model of delayed wound healing.

The proposed isolation protocol constitutes a significant improvement compared to UC, the gold-standard in the field. Isolated sEV maintain their regenerative properties, whereas downstream contaminants are minimized. The use of UF/SEC allows for the standardization and up-scalability required for mass production of sEV to be used in a clinical setting.

## 1 INTRODUCTION

Extracellular vesicles (EV) are a heterogeneous group of biological carrier systems that are released from most, if not all, cells. According to their size and origin, EV can be referred to as exosomes or microvesicles, if produced through the endosomal pathway or from budding of the plasma membrane, respectively. While exosomes range from 30-100 nm in diameter, microvesicles can measure up to 1000 µm (1,2).

EV are known to play a pivotal role as mediators of the communication between different cell types, namely through modulation and transport of RNAs (including microRNAs) (3,4). Apart from RNA, these vesicles carry a wide array of bioactive molecules, such as proteins and DNA, and have been demonstrated to be potential candidates for replacement of cell therapies in different disease contexts (5,6). Mesenchymal stem cells (MSCs) and mononuclear cells (MNC) obtained from umbilical cord blood (UCB) are among the most promising regenerative cell types (7,8) and constitute a rich source of EV (9). Current research efforts focus on exploring the potential of UCB-derived EV for the treatment of pathologies such as chronic wounds (10).

While EV have been extensively used in laboratory settings, their application to the clinics relies on the standardization of isolation and purification methods. Differential centrifugation, currently considered the gold-standard, leads to a significant retention of contaminants, such as soluble proteins (11). Moreover, this technique is highly influenced by human manipulation, time-consuming and includes multiple steps, that may result in sample loss (12,13).

In this study, we describe a scalable and clinically-compatible process of manufacturing UCB-MNC-derived EV. The combination of ultrafiltration and size exclusion chromatography (UF/SEC) reduces production time, while improving EV yield and standardization (14,15). Furthermore, we demonstrate the regenerative potential of our product, which significantly accelerates wound healing in a diabetic mouse model.

## 2 MATERIALS AND METHODS

### 2.1 Umbilical cord blood collection and cell culture

Human UCB samples were obtained upon signed informed consent, in compliance with Portuguese legislation. Donations were approved by the ethical committees of *Centro Hospitalar e Universitário de Coimbra*, Portugal. Samples were stored and transported to the laboratory in sterile bags with anticoagulant solution (citrate-phosphate-dextrose) and processed within 48 hours after collection.

2 million UCB-MNC/mL were cultured in X-VIVO 15 serum-free cell-culture medium (Lonza Group Ltd, Basel, Switzerland), supplemented with 0.5 μg/mL of FMS-like tyrosine kinase-3 and 0.5 μg/mL of stem-cell factor, under ischemia (0.5% O2). After 18 hours, conditioned media (CM) was collected for sEV purification.

### 2.2 UCB-MNC-sEV isolation

When indicated, sEV were purified by differential centrifugation as described in Thery C. et al. (12) (Figure 1A). For UF/SEC isolation (Figure 1B), CM was cleared by two sequential centrifugation steps at 300 xg (10 min) and 2000 xg (20 min), followed by filtration with 0.45 µm and 0.22 µm filters. A final ultrafiltration step was performed using VivaCell 250® (Sartorius AG, Goettingen, Germany), at 3 bar, through a 100 kDa filter. Small- (Superose 6 10/300 GL) or large-scale (Superose 6 pre-packed XK 26/70) SEC columns (GE Healthcare, Chicago, IL) were run with 500 µL or 15 mL of supernatant, at a flow rate of 0.5 mL/min or 3 mL/min, respectively. UV absorbance was measured at 220, 260 and 280 nm.

**Figure 1:**
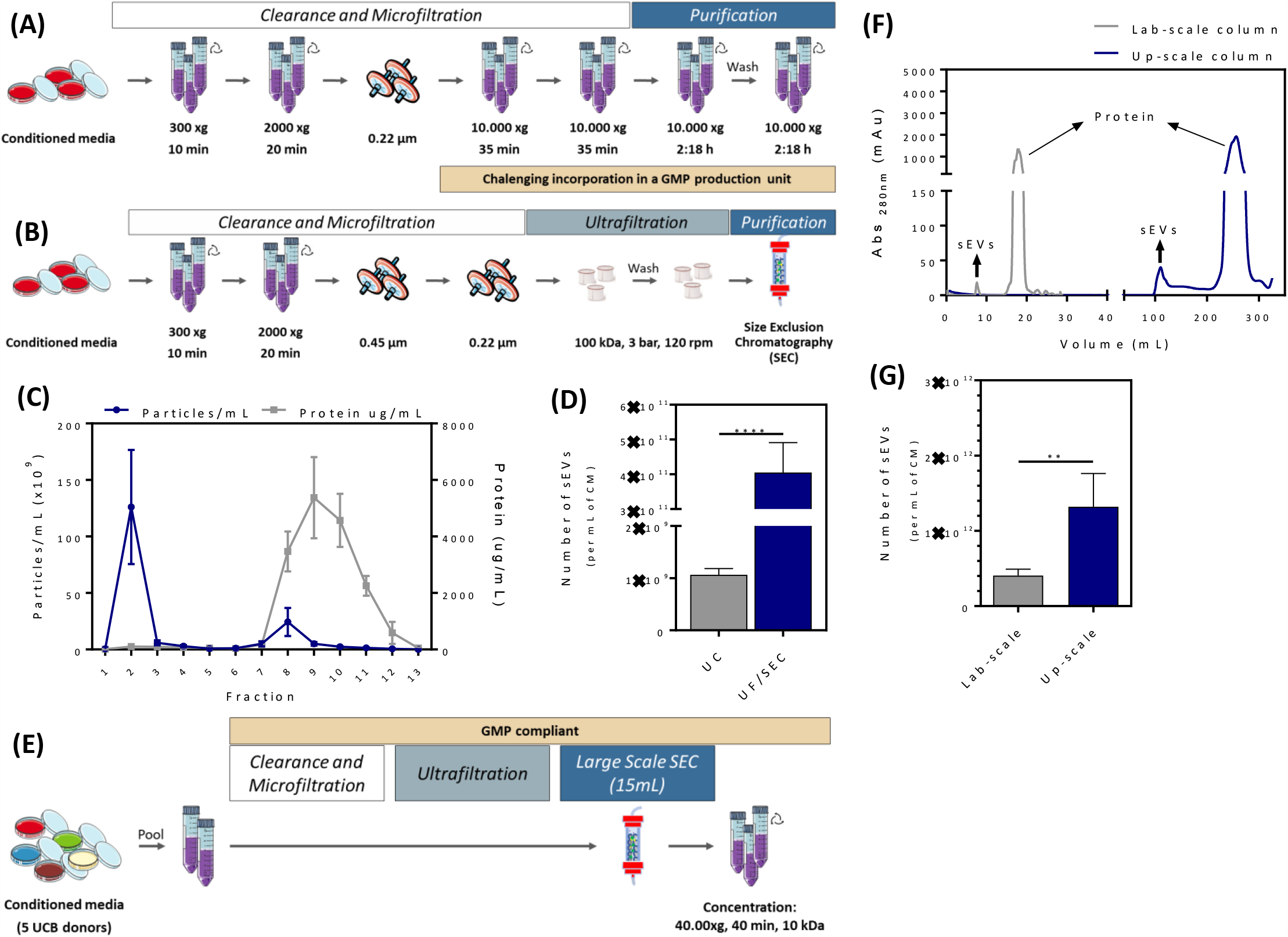
Comparison of ultracentrifugation (UC) and combined ultrafiltration and size exclusion chromatography (UF/SEC) for the isolation of small extracellular vesicles (sEV). Workflow for **(A)** UC and **(B)** UF/SEC. **(C)** Particle and protein concentration as a function of UF/SEC fraction. **(D)** sEV yield per mL of conditioned media (CM), using UC or UF/SEC (n = 5). **(E)** Workflow for up-scaled UF/SEC, based in the pooling from several UCB donors. **(F)** FPLC chromatogram comparing smaller and larger SEC columns. **(G)** sEV yield per mL of conditioned media (CM), using smaller- or larger-scale UF/SEC (n ≥ 7). All values are mean ± SEM. **p<0.01, ****p<0.0001

**Figure 2:**
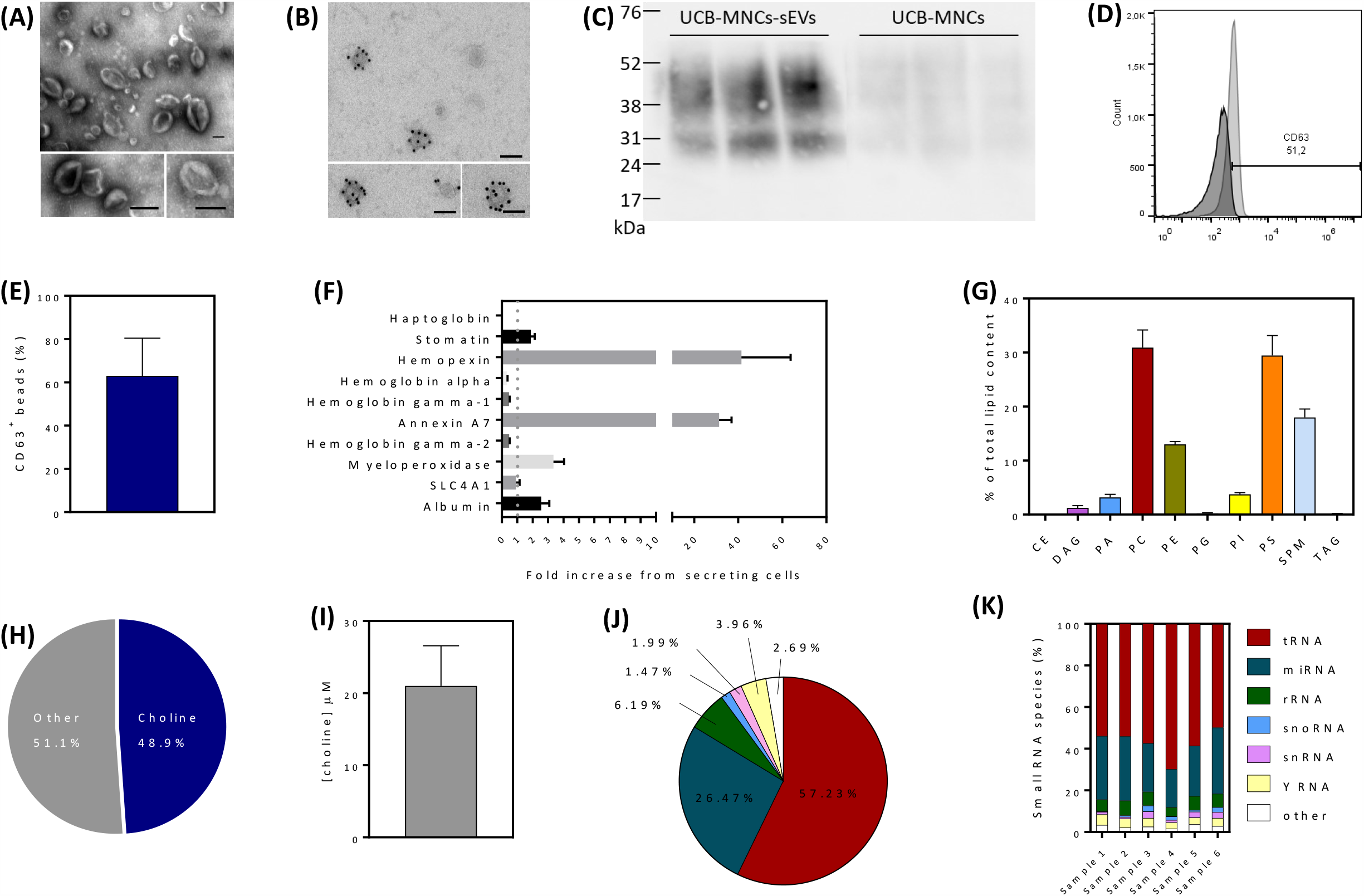
Characterization of small extracellular vesicles (sEV) isolated by an up-scaled protocol combining ultrafiltration and size exclusion chromatography (UF/SEC). TEM images of different samples isolated as described in Figure 1E, **(A)** unlabeled and **(B)** CD63-labelled. Scale bars = 100 nm. **(C)** Comparison of the CD63 content in sEV or their parent cells (umbilical cord blood-derived mononuclear cells, UCB-MNC), by western blot. **(D, E)** Flow cytometry analysis of CD63 on the surface of sEV (n = 3). Dark histogram corresponds to the unlabeled sample. **(F)** Fold increase of the proteins in sEV, compared to their parent cells, as identified by mass spectrometry (n = 3). **(G, H)** Mass spectrometry identification of lipids in sEV (n = 6). CE = cholesterol ester, DAG = diacylglycerol, PA = phosphatidic acid, PC = phosphatidylcholine, PE = phosphatidylethanolamine, PG = phosphatidylglycerol, PI = phosphatidylinositol, PS = phosphatidylserine, SPM = sphingomyelin, TAG = sphingomyelin. **(I)** Concentration of lipids with choline groups in sEV (n = 22). **(J, K)** Characterization of the RNA species in sEV by RNA-seq. All values are mean ± SEM.

### 2.3 Transmission Electron Microscopy (TEM)

Samples were mounted on 300 mesh formvar copper grids, stained with uranyl acetate 1 %, and examined with a Jeol JEM 1400 transmission electron microscope (Tokyo, Japan). Images were digitally recorded using a Gatan SC 1000 ORIUS CCD camera (Warrendale, PA), and photomontages were performed with Adobe Photoshop CS software (Adobe Systems, San Jose, CA), at the Institute for Molecular and Cell Biology (IBMC) of the University of Porto, Portugal.

For CD63 immunogold labelling, sEV were prepared according to Thery C. et al. (12). sEV were fixed with 2 % PFA and underwent single immunogold labeling with protein A conjugated to gold particles 10 nm or 15 nm in diameter (Cell Microscopy Center, Department of Cell Biology, Utrecht University, Netherlands). Grids were analyzed on a Tecnai Spirit G2 electron microscope (FEI, Eindhoven, The Netherlands) and digital acquisitions were made with a 4k CCD camera (Quemesa, Olympus, Muenster, Germany).

### 2.4 Flow cytometry

Briefly, 1×10^10^ sEV were incubated with 3.8 µm aldehyde/sulfate latex 4% (m/v) beads (Molecular probes, Eugene, OR). Coated beads were then incubated with anti-human CD63 (eBioscience, San Diego, CA) and analysed with a BD AccuriTM C6 (BD Biosciences, San Jose, CA).

### 2.5 Mass Spectrometry: Proteins

Tryptic peptides of purified sEV aliquots were analysed by liquid chromatography-mass spectrometry (LC-MS). For protein identification, an information-dependent acquisition (IDA) analysis by NanoLC-MS using TripleTOF 6600 (ABSciex, Framingham, MA) was used. Peptides were separated through reversed-phase chromatography (RP-LC) in a trap- and-elute mode. Trapping was performed at 2 µl/min with 0.1% formic acid, for 10 min, on a Nano cHiPLC Trap column (Sciex 200 µm × 0.5 mm, ChromXP C18-CL, 3 µm, 120 Å).

Separation was performed on a Nano cHiPLC column (Sciex 75 µm × 15 cm, ChromXP C18-CL, 3 µm, 120 Å) at a flow rate of 300 μL/min, applying a 90 min linear gradient of 5 % to 30 % (v/v) of 0.1% formic acid in acetonitrile.

Peptides were sprayed into the MS through an uncoated fused-silica PicoTip(tm) emitter (360 µm O.D., 20 µm I.D., 10 ± 1.0 µm tip I.D., New Objective). The source parameters were set as follows: 12 GS1, 0 GS2, 30 CUR, 2.4 keV ISVF and 100 °C IHT. An information dependent acquisition (IDA) method was set with a TOF-MS survey scan (400-2000 m/z) for 250 msec. The 50 most intense precursors were selected for subsequent fragmentation and the MS/MS were acquired in high sensitivity mode (150–1800 m/z for 40 ms each) with a total cycle time of 2.3 sec. The selection criteria for parent ions included a charge state between +2 and +5, and count above a minimum threshold of 125 counts per second. Ions were excluded from further MS/MS analysis for 12 sec. Fragmentation was performed using rolling collision energy with a collision energy spread of 5.

The obtained spectra were processed and analyzed using ProteinPilot(tm) software, with the Paragon search engine (version 5.0, Sciex). The following search parameters were set: search against Uniprot/SwissProt reviewed database restricted to Homo sapiens (accessed in May 2017); Iodoacetamide, as Cys alkylation; Tryspsin, as digestion; TripleTOF 6600, as the Instrument; ID focus as biological modifications and Amino acid substitutions; search effort as thorough; and a FDR analysis. Only the proteins with Unused Protein Score above 1.3 and 95% confidence were considered.

Data provided/obtained by the UniMS – Mass Spectrometry Unit, ITQB/IBET, Oeiras, Portugal. The obtained data was analyzed by functional enrichment analysis tool Funrich V3.0.

### 2.6 Mass Spectrometry: Lipids

Lipids were extracted using chloroform and methanol and samples spiked with specific internal standards prior to extraction. The mass spectra were acquired on a hybrid quadrupole/Orbitrap mass spectrometer equipped with an automated nano-flow electrospray ion source in both positive and negative mode. The identification of lipids was performed using LipotypeXplorer on the raw mass spectra. For MS-only mode, lipid identification was based on the molecular masses of the intact molecules. MS/MS mode included the collision induced fragmentation of lipid molecules and lipid identification was based on both the intact masses and the masses of the fragments.

### 2.7 RNAseq

Total RNA was isolated using Exiqon miRCURY isolation kit (Qiagen, Hilden, Germany) and small RNA quality and quantification was performed in Bioanalyzer 2100 (Agilent Technologies, Santa Clara, CA). To construct each library, 1.5 ng of small RNA were hybridized and ligated to Ion Adapters v2 (Life Technologies, Carlsbad, CA). Following reverse transcription, purified cDNA samples were size-selected, amplified by PCR and further purified. cDNA samples were barcoded using Platinum PCR SuperMix High Fidelity polymerase and Ion Xpress RNA-Seq Barcode 1–16 Kit (Life Technologies, Carlsbad, CA). Yield and size distribution of the cDNA libraries were assessed using a DNA1000 chip (Agilent Technologies, Santa Clara, CA). Total barcoded cDNA within the 50–300 base pair range was considered to be derived from small RNA. 50 picomoles of each barcoded library were pooled and clonally amplified onto Ion Sphere Particles (ISPs), enriched and loaded in an Ion 530 Chip (Life Technologies, Carlsbad, CA). Enriched ISPs were sequenced using Ion S5 (Life Technologies, Carlsbad, CA) with 500 flows. Small RNA sequencing was performed at GenCore (i3s, Porto, Portugal).

Small-RNA sequencing data was quality controlled with FASTQC. No hard sample trimming was performed, and the aligner’s own ability to ignore regions in reads which did not map to the human genome was used. Alignment was performed using aligner Bowtie2 and samples were aligned to the RefSeq genome GRCh38 (GCF_000001405.26_GRCh38_genomic), as it matches the genome used by the database MIRBASE for identification and annotation of miRNAs. Cufflinks tool was used for annotation and estimation of the relative abundance of each gene. Cufflinks was performed using annotation against the whole human genome by RefSeq, to detect all small RNAs. The obtained data was generated by Bioinf2Bio (Porto, Portugal).

### 2.8 Matrigel tube formation assay

10,000 endothelial cells/well were seeded on polymerized Matrigel (BD Biosciences, San Jose, CA) in a 96-well plate in complete endothelial media (ATCC, Manassas, VA). After 24h, the media was replaced with a starvation media, either alone or supplemented with 1×10^10^ sEV/mL. Cells were photographed, after 4 and 8 hours, using an InCell microscope (GE Healthcare, Chicago, IL) (Figure 3A). The number of nodes and meshes, total length and total segment length was quantified with ImageJ (macro angiogenesis) (NIH, Bethesda, MD).

**Figure 3:**
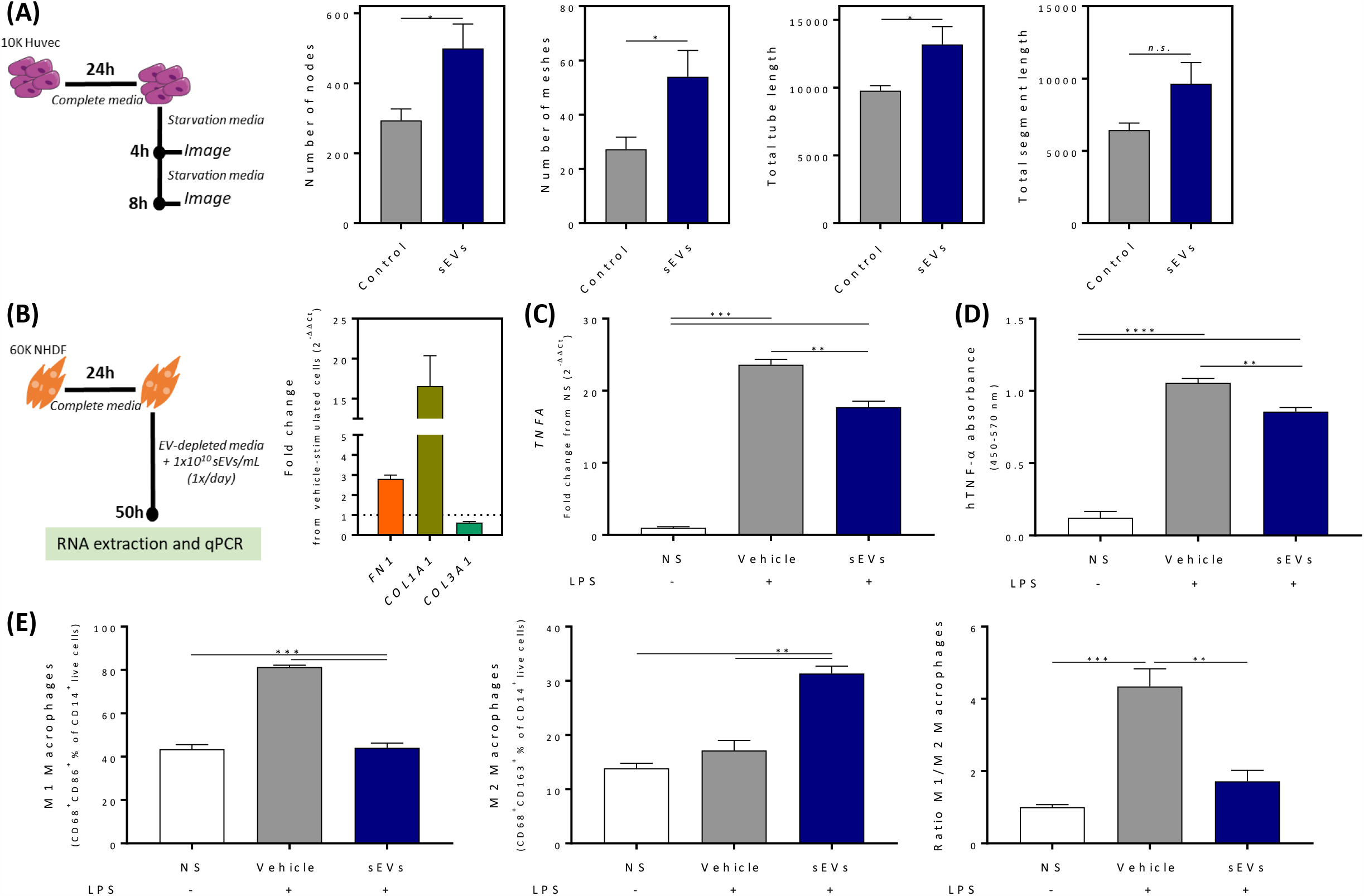
Contribution of UCB-MNC-sEV towards angiogenesis, collagen production and macrophage phenotype, *in vitro*. **(A)** Matrigel tube formation assay, using human umbilical vein endothelial cells (HUVEC), incubated or without 1×10^9^ sEV/mL. Number of nodes, meshes, total length and segment length were evaluated by ImageJ (n = 3). **(B)** Fibronectin (*FN1*), collagen I (*Col1A1*) and collagen III (*Col3A1*) expression in normal human dermal fibroblasts (NHDF), by RT-qPCR (n = 3), with or without treatment of 1×10^9^ sEV/ml. **(C, D)** TNFα expression and release by THP-1-derived macrophages (n = 8), determined by RT-qPCR and ELISA, respectively, with or without treatment of 1×10^10^ sEV/ml **(E)** Effect of sEV on the phenotype of LPS-stimulated macrophages (n = 8). THP-1 cells were differentiated into macrophages with PMA (25nM), before stimulation with LPS (1 µg/mL), and cells were incubated with or without 1×10^10^ sEV/ml. All values are mean ± SEM. *n*.*s*. = non-significative, *p<0.05, **p<0.01, ***p<0.001, ****p<0.0001

### 2.9 Expression of ECM components

60,000 normal human dermal fibroblasts (NDHF) were seeded in a 12 well plate in ATCC fibroblast supplemented medium for 24h, after which the medium was replaced by sEV-depleted medium (Figure 3B). Treatment with 1×10^10^ sEV/mL was administrated daily for 3 days (without replacement of medium). Control cells were treated with filtered PBS. mRNA analysis of ECM genes was performed by qPCR with the primers listed in Table 1. Beta-actin (ACTB) was used as a house-keeping gene. Data was analysed with CFX Manager software (Bio-Rad, Hercules, CA).

**Table 1.**
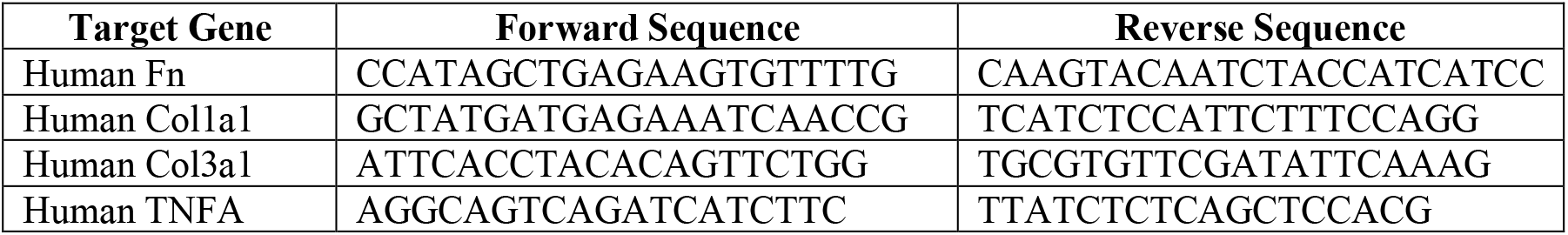
Primer sequences for qPCR-amplified genes.

### 2.10 Macrophages assays

THP-1 cells were differentiated into macrophages with 25 mM PMA, for 72 h. After stimulation with 1µg/mL LPS, cells were treated with 1×10^10^ sEV/mL for 24 h. Levels of TNF-α mRNA and protein were detected by qPCR (Table 1) and ELISA (BioLegend, San Diego, CA), respectively.

For flow cytometry, macrophages were stained with fluorescently labelled anti-human CD14 (M5E2), CD68 (Y1/82A), CD86 (FUN-1) and CD163 (GHI/61), all from BD Biosciences (San Jose, CA). M1 macrophages were defined as CD14^-^CD68^+^CD86^+^ and M2 macrophages as CD14^-^CD68^+^CD163^+^.

### 2.11 *In vivo* wound healing

Animal testing protocols were approved by the Portuguese National Authority for Animal Health (DGAV) and performed respecting national and international animal welfare regulations. Male C57BL/6 wild-type mice (8-10 weeks-old), purchased from Charles River (Écully, France), were housed in a conventional animal facility on a 12-hr light/12-hr dark cycle and fed regular chow *ad libitum*.

Diabetes mellitus was induced by daily intraperitoneal injection of 50 mg/kg streptozotocin (STZ, Sigma-Aldrich, St. Louis, MO), for 5 consecutive days (Figure 4A). Glycemia was monitored weekly (Accu-Chek Aviva, Roche, Basel, Switzerland) and only animals with blood glucose levels higher than 300 mg/dL were used. The animals were diabetic for 6-8 weeks prior to wound induction. When necessary for weight maintenance, 16-32 U/kg of insulin (Sigma-Aldrich, St. Louis, MO) were injected subcutaneously.

**Figure 4:**
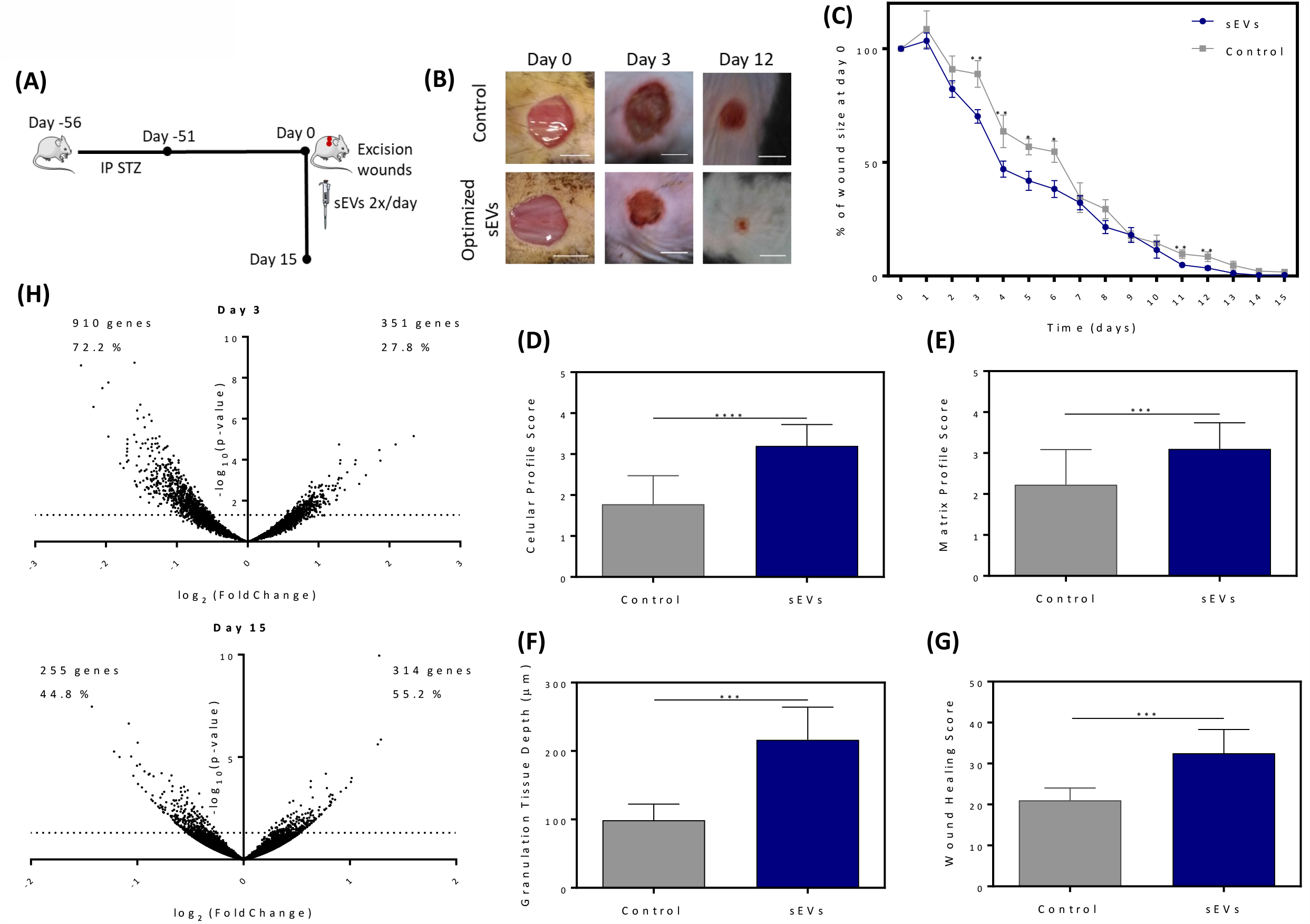
Effect of small extracellular vesicles (sEV) in wound healing. **(A)** Workflow of *in vivo* experiment, using a streptozotocin (STZ)-induced diabetic mouse model. sEV (at 2.5×10^9^ sEV/ml) or PBS (control) were administered topically twice per day, after performing two excisional wounds on the back of mice. **(B)** Representative wound micrographs of each condition at days 0, 3 and 12. Scale bar = 0.5 cm. **(C)** Percentage of wound size at day 0, measured daily during 15 days (n ≥ 4). **(D-G)** Scoring of the cellular and matrix profiles, granulation tissue depth and wound healing (n ≥ 3). Profiles were scored as follows: cellular (0 = low/minimal cellular response, 1 = predominantly inflammatory, 2 = mixed inflammatory and proliferative, 3 = predominantly proliferative, 4 = entirely proliferative); matrix (0 = no/minimal new matrix, 1 = predominantly fibrinous, 2 = mixed fibrinous and collagenous, 3 = predominantly collagenous, 4 = entirely collagenous); wound healing score is based on cellular and matrix profiles. **(H)** Volcano plot displaying differentially expressed genes between control and sEV-treated wounds at days 3 and 15 (n = 3). Dotted line represents a p-value of 0.05. All values are mean ± SEM. *p<0.05, **p<0.01, ***p<0.001, ****p<0.0001

For wound induction, animals were anesthetized by intramuscular injection of a xylazine/ketamine solution (ketamine hydrochloride, 10 mg/mL, (Imalgene®, Merial, Barcelona, Spain), 50 mg/kg of body weight, and xylazine hydrochloride, 2 mg/mL, (Rompun®, Bayer Healthcare, Germany), 10 mg/kg of body weight), and the dorsal trunk was shaved and disinfected with a povidone-iodine solution. Two 6 mm diameter full-thickness excision wounds were performed with a sterile biopsy punch in the dorsum of each animal.

Treatments (2.5×10^8^ sEV or PBS) were applied immediately after wound excision and repeated as mentioned in Figure 4A (2x/day for 15 days). Dose was chosen in accordance with previous experiments (10). Animals were monitored daily for the total duration of the experiment. Individual wounds were traced onto acetate paper and wound size was determined with ImageJ (NIH, Bethesda, MD).

Total RNA from murine skin biopsies was obtained using conventional Trizol extraction (NZYTech, Lisbon, Portugal). cDNA libraries for each sample were generated at Gene Expression Unit (Instituto Gulbenkian de Ciência, Oeiras, Portugal) from 50 ng of total RNA, using Quantseq 3′ mRNA-Seq Library Prep kit (Lexogen, Vienna, Austria). Data was analyzed using QuantSeq data analysis pipeline on the Bluebee genomic platform (Bluebee, Rijswijk, Netherlands). Funrich and Ingenuity Pathways Analysis software were used for the identification of biological processes and pathways affected in response sEV. Only differentially expressed genes with a p-value smaller than 0.05 and count number higher than 10 were considered.

A second *in vivo* experiment was performed by Cica Biomedical (Knaresborough, UK). Mice (BKS.Cg-m Dock7m +/+ Leprdb /J, CRL. Italy) were anaesthetised using isofluorane and air; and their dorsal flank skin was clipped and cleansed. A single standardised full-thickness wound (10mm × 10mm) was created on the left flank approximately 10mm from the spine.

All wounds were then dressed with the transparent film dressing Tegaderm™ Film (3M Deutschland GmbH, Germany); after which they topically received one of the treatments (2.5×10^8^ sEV or PBS) described below applied by injection through the Tegaderm™ film using a 27-gauge needle. All animals were terminated on post-wounding day 20. Harvested tissues were fixed and embedded in paraffin wax. Paraffin embedded wounds were then sectioned (4-6μm) and representative sections (from the centre of each wound) stained with Haematoxylin & Eosin - to facilitate measurement of granulation tissue depth, % cranio-caudal contraction and wound healing progression. Skin condition was scored blindly based on histological data obtain by an independent pathologist, as follows: cellular profile (0 = low/minimal cellular response, 1 = predominantly inflammatory, 2 = mixed inflammatory and proliferative, 3 = predominantly proliferative, 4 = entirely proliferative); matrix profile (0 = no/minimal new matrix, 1 = predominantly fibrinous, 2 = mixed fibrinous and collagenous, 3 = predominantly collagenous, 4 = entirely collagenous); wound healing score is based cellular and matrix profiles.

### 2.12 Statistical Analysis

Statistical analyses were performed by student t-test or by a one or two-way ANOVA test followed by a Bonferroni’s multiple comparisons test. Statistical analyses were performed by GraphPad Prism 6 software (GraphPad Software, Inc.). Significance levels were set at *P < 0.05, **P < 0.01, ***P < 0.001, and ****P < 0.0001.

## 3 RESULTS

### 3.1 Optimization and up-scaling of small extracellular vesicle (sEV) purification

Although UC (Figure 1A) is viewed as the gold-standard for EV isolation (16), UF/SEC (Figure 1B) has been recently described to be considerably faster and have less variability (14). Using the latter method, small EV (sEV, with a size range of 50 to 200 nm) can be detected by SEC and eluted within the void volume (Figure 1C), accounting for approximately 80% of its content (Figure S1A,C). A second peak, corresponding to approximately 60 kDa, is consistent with the presence of albumin, a frequent contaminant of sEV samples from biological fluids (11) (Figure S2).

With the goal of establishing an optimized protocol, sEV were isolated using either UC or UF/SEC. By directly comparing both methods, no differences were found in the population distribution of isolated particles (Figure S1A). However, UF/SEC allowed for a significantly higher yield of sEV per volume of CM (roughly 400 times higher) than traditional UC (Figure 1D).

Considering the upcoming use of EV for clinical application, which requires standardized mass production, we up-scaled the above-mentioned UF/SEC protocol, to enable future compliance with GMP standards (Figure 1E). No differences regarding particle population or modal size were detected between the lab-scale or up-scale protocols (Figure 1F and S1C,D). Remarkably, using a larger SEC column more than doubled sEV yield per volume (Figure 1G), further substantiating the advantages of this protocol. Thus, sEV can be efficiently isolated using an up-scaled method combining UF and SEC.

### 3.2 sEV characterization

sEV isolated with our optimized method were extensively characterized, regarding their protein, lipid and RNA content. These vesicles have the expected size and morphology of sEV (Figure 2A) and are enriched in CD63 (Figure 2B-E), a classical marker of exosomes, suggesting an endocytic origin (17). Further characterization, using LC-MS, revealed various proteins associated with sEV (Table S1), several of which were shown to be more abundant in sEV relative to their parent cell (Figure 2F), a finding which may imply active sorting of particular proteins into sEV. Importantly, a bioinformatic analysis using Funrich® unveiled a significant association between proteins found in sEV and biological pathways involved in wound healing (Table S2).

For the identification of the major lipid species present in sEV, shotgun MS was employed (Figure 2G). While sphingomyelin (SPM) and phosphatidylethanolamine (PE) represent roughly 30 % of total lipids, phosphatidylcholines (PC) and phosphatidylserines (PS) account for nearly 60 % of lipid content. Additionally, the amount of cholesterol-fatty acid ester (CE) and triacylglyceride (TAG) lipids in sEV preparations is negligible, indicating that lipoproteins and lipid droplets are not co-isolated with sEV. Given that phospholipids containing a choline group (SPM and PC) represent almost 50 % of sEV lipids (Figure 2H), commercially available choline kits can be used for routine quality control. sEV purified with this up-scaled UF/SEC protocol contain an average choline concentration of 21 µM (Figure 2I). Of note, the lipids present in the purified UCB-MNC-sEV are composed of very long fatty acid chains (VLFAC), such as SPM (42:1 and 42:2), PS (38:4) and PC (34:1) (Figure S4). This is in accordance with what has been described for sEV/exosomes isolated from other cells (18), indicating that there is a specific sorting of VLFAC lipids to sEV.

The genetic cargo of isolated sEV was characterized via RNA sequencing. In all analyzed samples, tRNAs (57 %) and miRNAs (27 %) correspond to a total of 84 % of the RNA cargo (Figure 2J,K). From a total of 175 miRNAs identified in 6 sEV samples, 35 (20 %) were common to all samples (Table S3 and Figure S5), among which mir-150 and mir-223 are the most abundant and have been described as beneficial during wound healing (10,19).

Therefore, based on the described characteristics, we suggest that sEV quality control thresholds can be defined based on the concentration of: i) nanoparticles; ii) total proteins; iii) albumin; iv) total lipids; v) choline phospholipids; vi) total RNAs; and vii) specific small RNAs.

### 3.3 Bioactivity of sEV *in vitro*

In a previous work, UCB-MNC-sEV isolated by UC were shown to accelerate wound healing *in vivo* (10). In order to validate if their bioactivity was maintained when isolated with an optimized and up-scaled GMP-compatible method, endothelial cells (HUVEC) or fibroblasts (NHDF) were stimulated *in vitro* with isolated vesicles. As shown in Figure 3A and Figure S8, sEV promote angiogenesis, demonstrated by the increase in the number of nodes, meshes and tube length formed by HUVEC (Figure S8). Furthermore, after stimulation with sEV, NHDF increase the expression of extracellular matrix (ECM) proteins important during wound healing, such as collagens and fibronectin (Figure 3B). Moreover, sEV significantly reduce the expression and release of TNFα by LPS-stimulated macrophages (Figure 3C,D), a phenomenon that is likely due to a shift from the pro-inflammatory M1 to the anti-inflammatory M2 phenotype (Figure 3E). Altogether, these results suggest that UCB-MNC-sEV have potential beneficial effects during wound healing, as well as anti-inflammatory properties.

### 3.4 Bioactivity of sEV *in vivo*

Having established a significant effect of sEV isolated in large scale, using UF/SEC, we next addressed the bioactivity of these vesicles in an *in vivo* model of delayed wound healing (Figure 4A). Macroscopically, sEV significantly accelerated wound closure, particularly in the initial phase of healing (Figure 4B,C). After 12 days of treatment, control-treated wounds were still significantly larger than wounds treated with sEV, proving a beneficial effect of the vesicles also at later stages of healing.

Based on the characteristics of the cells found in each wound (Figure S9), as well as on the quality of newly-formed matrix, mice were scored according to their cellular and matrix profiles, respectively. While control-treated animals displayed a mixed proliferative and inflammatory cellular score at the end of the experiment, sEV-treated wounds had a predominantly proliferative profile, with few or no inflammatory cells (Figure 4D).

Additionally, sEV treatment resulted in a predominantly collagenous matrix, whereas control-treated wounds had a mixed fibrinous and collagenous matrix profile (Figure 4E). Consistently, granulation tissue, i.e. new connective and endothelial tissue, was significantly thicker in sEV-treated animals (Figure 4F). These histological data show that, at the time of sacrifice, sEV-treated wounds were in a more advanced stage of healing than control wounds, a finding that is summarized in Figure 4G. Thus, while macroscopic differences are mostly seen in the six initial days of treatment, histological analyses show that sEV significantly improve wound healing at every stage.

### 3.5 Genetic signature of sEV-treated wounds

In order to unveil the mechanisms involved in sEV-triggered wound healing, global gene expression analysis was performed at days 3 and 15. These time-points were chosen according to literature, corresponding to inflammatory and remodelling wound healing phases, respectively (20,21). When comparing sEV-treated wounds with control wounds, we found that 1261 genes (19.9 %) are differentially expressed at day 3 (p<0.05). Of these, the large majority (72 %) is down-regulated in sEV-treated wounds (Figure 4H). Down-regulated genes at day 3 are likely related with inflammatory processes, since iNOS, IL-6, TNFα and CXCL1 were found to be significantly or tendentially down-regulated in sEV-treated samples (Figure S6A-D). Accordingly, CD163 and arginase-1, two molecules associated with anti-inflammatory M2 macrophages, were found to be up-regulated in wounds treated with sEV for 3 days (Figure S6E,F). At day 15, 569 genes (6 %) were found to be differentially expressed, of with approximately half was down-regulated (Figure 4H). Funrich analysis showed that biological processes related with inflammatory and apoptotic responses are more representative at day 3, while cell proliferation, cell adhesion and ECM organization have a heavier impact at day 15 (Table S4). These findings suggest that UCB-MNC-sEV have an initial anti-inflammatory effect, which is followed by an impact in tissue remodeling during skin wound healing.

## 4 DISCUSSION

EV from UCB have the potential to become a powerful tool in regenerative medicine (10,22). As such, recent efforts have focused on optimizing the isolation process of these vesicles (23). Herein, we describe an optimized and up-scaled isolation process for sEV, using UF/SEC, that allows for a higher yield and lower contamination with EV larger than 200 nm (sEV), compared with traditional UC, considered the gold-standard in EV isolation. Using UF/SEC, we were able to recover approximately 400 times more sEV per mL of media than with UC, and up-scaling this process further increases EV yield by about 3-fold. The increased yield per mL of CM results in an overall lower contamination of the vesicles with serum proteins, such as albumin (Figure S3), the major contaminant in sEV purified from biological materials (24,25). These results are in line with prior observation when using UF/SEC methodology to isolate sEV (26–28). Depending on the specification of each described method, UF/SEC lead to no yield or 5-fold yield in comparison to UC while sEV biophysical characteristics were conserved. Nevertheless, our methodology was design to overcome a common issue in sEV isolation, the initial volume of biofluids or cell culture medium. Firstly, we scaled up our process which is able to deal with large initial volumes that after concentration by ultrafiltration is read to be injected in a SEC column. In this regard, we also up scaled from a small to a large-scale SEC column which allows up to 15 mL of supernatant injection. Of note, this SEC column can be even replaced by larger ones compatible with GMP production facilities. In addition, the implemented SEC columns are suitable for biofluids injection as their matrix effectively resolves EVs and high-density lipoprotein in contrast with others (26).

Characterizing the content of sEV is essential not only to understand their mechanism of action, but also to be able to control and standardize isolated vesicle batches. In line with previous literature (29–32), sEV isolated with the described protocol were shown to be enriched in proteins like hemopexin, annexin A7 and myeloperoxidase (MPO). Hemopexin is a heme-binding plasma protein, described as a potent anti-inflammatory agent preventing heme-toxicity (33), whereas MPO is a heme-containing peroxidase with antimicrobial activity (34). Therefore, it is plausible that both proteins contribute to wound healing by reducing inflammation, while preventing bacterial growth, two processes that are known to accelerate wound healing. In fact, preliminary *in vitro* data (not shown) demonstrate that UCB-MNC-sEV can inhibit the growth of bacterial strains found in human skin, a finding that is supported by previous literature (35).

Similarly to sEV isolated from other cell sources (18), phosphatidylcholines, phosphatidylserines and sphingomyelin are the three major phospholipid constituents of UCB-MNC-sEV. These results reinforce previous literature describing sEV as having a higher membrane rigidity than microvesicles and apoptotic bodies (36). Aside from being an important structural component, lipids also play a key role in the interaction of sEV with recipient cells (37). Nevertheless, their specific role in wound healing remains poorly explored. Sphingolipids, which comprise sphingomyelins, have been identified as important wound healing mediators, namely through the modulation of inflammation (38,39). Also, sphingosine-1-phosphate promoted keratinocyte migration (40) whereas PE induced an antifibrotic phenotype in human fibroblasts (41). Moreover, PC-liposomes treatment displayed wound-healing and anti-inflammatory properties in a guinea pig model (42). These evidences reinforce sEV lipid composition as a crucial characteristic for their bioactivity in tissue regeneration. A major concern regarding small vesicle purification is their contamination with lipid droplets or lipoproteins. Due to their biophysical characteristics, similar to sEV, these cannot easily be eliminated (43). In contrast to what has been described for UC (44), sEV isolated with UF/SEC have a low content in CE and TAG. Overall, our results indicate that this optimized methodology is better suited for future clinical applications than UC, since it results in sEV with lower protein and lipid contamination.

Small RNAs found in sEV are known to have therapeutic benefits, particularly in wound healing. The described protocol allows for the isolation of sEV rich in microRNA with a well-documented role in skin regeneration: miR-150 and miR-205 improve keratinocyte and fibroblast function (10,45), miR-146a targets TNF-alpha during inflammation(46), and miR-221 modulates the angiogenic activity of stem cell factor via its c-kit receptor (47).

Accordingly, UCB-MNC-sEV promote angiogenesis and expression of collagen 1, a major protein found in connective tissues. Furthermore, these vesicles were shown to be beneficial during wound healing in diabetic mice, in agreement with previous data obtained with UC-isolated sEV (10). The effect of sEV treatment was macroscopically discernible primarily in the first days of healing, and histological data obtained at the end of the experiment showed significantly more regeneration in wounds treated with sEV than in controls. The acceleration of wound closure in an initial phase may provide advantages for reduction of bacterial infections, common in wound patients, facilitating progression into a non-inflammatory remodeling phase.

Indeed, evidences of sEV’ anti-inflammatory function were abundant in our experiments. Genetic expression of diabetic wounds revealed the effect of sEV in genes related with inflammatory processes. Some of these genetic changes seem to affect the fate of macrophages, from an M1 (inflammatory) to an M2 (anti-inflammatory) phenotype. iNOS, an enzyme present in M1 macrophages, is down-regulated, while M2-associated proteins CD163 and arginase-1 are up-regulated in sEV-treated wounds. These observations are corroborated by *in vitro* data, showing the preferential shift towards M2 in sEV-stimulated macrophages. As chronic wounds are stalled in the inflammatory phase (48), an increase in anti-inflammatory macrophages is likely crucial for progression towards the remodeling phase of healing.

## 5 CONCLUSION

In summary, we observe that UCB-MNC-sEV isolated with the described optimized method are obtained with high purity and significantly increased yield, while enclosing a cocktail of proteins, lipids and RNAs which promote wound healing, by modulation of inflammation, angiogenesis and ECM remodeling (Figure S7). This optimized protocol can be easily adapted for mass production of pure and well-characterized vesicles in GMP facilities for clinical use. Furthermore, control and standardization of isolated vesicle batches will be ensured by the well-defined quality control attributes established in this work.

## ACKNOWLEDGMENTS

This work was co-funded by Regional Operational Program Center 2020, Portugal 2020 and European Union through FEDER within the scope of CENTRO-01-0247-FEDER-022398 and supported by the FCT fellowship SFRH/BD/137633/2018. The authors greatly appreciated the assistance of Graça Raposo for revising this manuscript and for helpful discussions.

## DISCLOSURE OF POTENTIAL CONFLICTS OF INTEREST

R.M.S.C and J.S.C are inventors of the patent PCT/IB2017/000412 (Use of umbilical cord blood derived exosomes for tissue repair) and R.M.S.C., S.C.R and J.S.C. are inventors of the patent PCT/IB2019/058462 (Compositions comprising small extracellular vesicles derived from umbilical cord blood mononuclear cells with anti-inflammatory and immunomodulatory properties), currently explored by Exogenus Therapeutics, S.A. Financial interest is claimed by Exogenus Therapeutics, S.A., which holds a licence (PCT/IB2017/000412) and a patent related to this work (PCT/IB2019/058462), and J.S.C. in the capacity of a shareholder of Exogenus Therapeutics, S.A. The other authors declare no conflicts of interest.

## DATA AVAILABILITY STATEMENT

For any data requests, please contact Exogenus Therapeutics, S.A., at.

## Abbreviation list

ACTB: beta-actin
CE: cholesterol-fatty acid ester
CM: conditioned medium
ECM: extracellular matrix
EV: Extracellular Vesicles
HUVEC: human umbilical vein endotelial cells
ISPs: Ion Sphere Particles
MNC: mononuclear cells
MSC: Mesenchymal stem cells
NDHF: human dermal fibroblasts
PC: phosphatidylcholines
PE: phosphatidylethanolamine
PS: phosphatidylserines
sEV: small Extracellular Vesicles
SPM: sphingomyelin
TAG: triacylglyceride
UC: ultracentrifugation
UCB: umbilical cord blood
UF/SEC: ultrafiltration combined with size exclusion chromatography
VLFAC: very long fatty acid chains

