## Supplementary Data for "Development of an optimized and scalable method for isolation of umbilical cord blood-derived small extracellular vesicles for future clinical use"

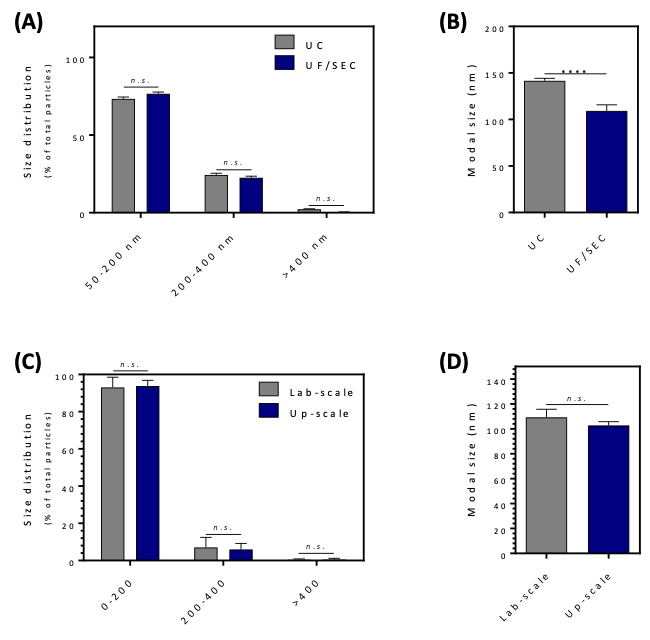

**Figure S1: Influence of isolation method on population and size characteristics of nanoparticles derived from umbilical cord blood mononuclear cells.** **(A)** Population distribution, by nanoparticle tracking analysis (NTA), of particles (sEV) isolated with ultracentrifugation (UC) versus ultrafiltration and size-exclusion chromatography (UF/SEC) (n ≥ 4). **(B)** sEV modal size (n ≥ 4). The significantly higher values seen with UC are likely due to a higher contamination with protein complexes, which can be easily eliminated with SEC. **(C)** NTA population distribution of UF/SEC-isolated sEV, using a small- or large-scale column (n ≥ 12). **(D)** Modal size of sEV isolated with a lab-scale versus up-scale column (n ≥ 12). All values are mean ± SEM. *n.s.* = non-significative, ****p<0.0001.

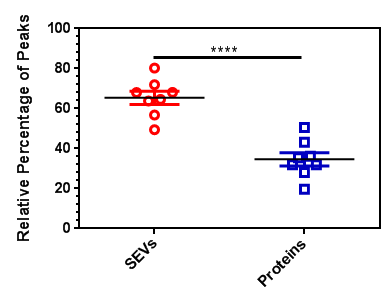
**Figure S2: Relative percentage of sEV and free protein in sEV samples isolated by UF/SEC.** Values are mean ± SEM. ****p<0.0001.

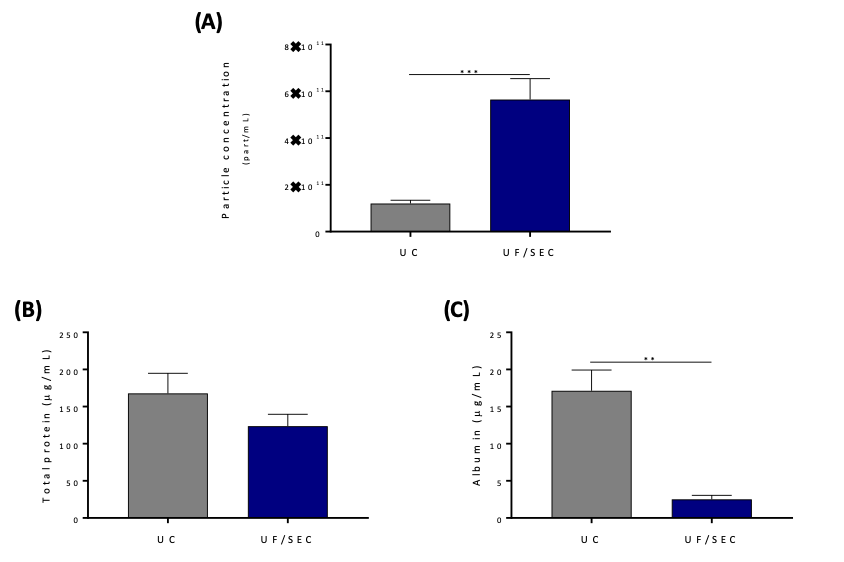

**Figure S3: Influence of isolation method on particle purity.** Concentration of **(A)** sEV (n ≥ 14), **(B)** total protein (n ≥ 17), and **(C)** albumin (n ≥ 4) in sEV samples isolated with UC versus UF/SEC. All values are mean ± SEM. **p<0.01, ***p<0.001.

**Table S1.** List of sEV-associated proteins, identified by LC-MS, in sEV samples isolated with UF/SEC (n = 3). Results were analyzed against ExoCarta data.

| **Ranking** | **UniProt ID** | **UniProt accession** | **Gene Symbol** | **Protein name** |
| --- | --- | --- | --- | --- |
| 2 | ALBU_HUMAN | P02768 | ALBU | Serum albumin |
| 10 | MYH9_HUMAN | P35579 | MYH9 | Myosin-9 |
| 13 | ANXA6_HUMAN | P08133 | ANXA6 | Annexin A6 |
| 17 | FLNA_HUMAN | P21333 | FLNA | Filamin-A |
| 23 | TFR1_HUMAN | P02786 | TFR1 | Transferrin receptor protein 1 |
| 32 | ANXA7_HUMAN | P20073 | ANXA7 | Annexin A7 |
| 34 | TLN1_HUMAN | Q9Y490 | TLN1 | Talin-1 |
| 36 | PDC6I_HUMAN | Q8WUM4 | PDC6I | Programmed cell death 6-interacting protein |
| 39 | AT1A1_HUMAN | P05023 | AT1A1 | Sodium/potassium-transporting ATPase subunit alpha-1 |
| 44 | HSP7C_HUMAN | P11142 | HSP7C | Heat shock cognate 71 kDa protein |
| 46 | FLOT1_HUMAN | O75955 | FLOT1 | Flotillin-1 |
| 47 | H4_HUMAN | P62805 | H4 | Histone H4 |
| 49 | TERA_HUMAN | P55072 | TERA | Transitional endoplasmic reticulum ATPase |
| 57 | CO3_HUMAN | P01024 | CO3 | Complement C3 |
| 59 | MOES_HUMAN | P26038 | MOES | Moesin |
| 63 | ANXA5_HUMAN | P08758 | ANXA5 | Annexin A5 |
| 69 | ANX11_HUMAN | P50995 | ANX11 | Annexin A11 |
| 70 | ANXA1_HUMAN | P04083 | ANXA1 | Annexin A1 |
| 79 | A2MG_HUMAN | P01023 | A2MG | Alpha-2-macroglobulin |
| 83 | CLH1_HUMAN | Q00610 | CLH1 | Clathrin heavy chain 1 |
| 89 | ITB1_HUMAN | P05556 | ITB1 | Integrin beta-1 |
| 87 | 4F2_HUMAN | P08195 | 4F2 | 4F2 cell-surface antigen heavy chain |
| 88 | G3P_HUMAN | P04406 | G3P | Glyceraldehyde-3-phosphate dehydrogenase |
| 92 | ANXA2_HUMAN | P07355 | ANXA2 | Annexin A2 |
| 94 | ENOA_HUMAN | P06733 | ENOA | Alpha-enolase |
| 100 | PRDX2_HUMAN | P32119 | PRDX2 | Peroxiredoxin-2 |
| 105 | SDCB1_HUMAN | O00560 | SDCB1 | Syntenin-1 |
| 104 | RAP1B_HUMAN | P61224 | RAP1B | Ras-related protein Rap-1b |
| 111 | KPYM_HUMAN | P14618 | KPYM | Pyruvate kinase PKM |
| 114 | HS90A_HUMAN | P07900 | HS90A | Heat shock protein HSP 90-alpha |
| 121 | 1433Z_HUMAN | P63104 | 1433Z | 14-3-3 protein zeta/delta |
| 128 | ACTN1_HUMAN | P12814 | ACTN1 | Alpha-actinin-1 |
| 131 | LDHA_HUMAN | P00338 | LDHA | L-lactate dehydrogenase A chain |
| 137 | PGK1_HUMAN | P00558 | PGK1 | Phosphoglycerate kinase 1 |
| 136 | GNAI2_HUMAN | P04899 | GNAI2 | Guanine nucleotide-binding protein G(i) subunit alpha-2 |
| 135 | BASI_HUMAN | P35613 | BASI | Basigin |
| 134 | ALDOA_HUMAN | P04075 | ALDOA | Fructose-bisphosphate aldolase A |
| 143 | PROF1_HUMAN | P07737 | PROF1 | Profilin-1 |
| 149 | GELS_HUMAN | P06396 | GELS | Gelsolin |
| 172 | 1433E_HUMAN | P62258 | 1433E | 14-3-3 protein epsilon |
| 180 | RAN_HUMAN | P62826 | RAN | GTP-binding nuclear protein Ran |
| 179 | PPIA_HUMAN | P62937 | PPIA | Peptidyl-prolyl cis-trans isomerase A |
| 186 | CD63_HUMAN | P08962 | CD63 | CD63 antigen |
| 187 | RAB7A_HUMAN | P51149 | RAB7A | Ras-related protein Rab-7a |
| 202 | CLIC1_HUMAN | O00299 | CLIC1 | Chloride intracellular channel protein 1 |
| 214 | LDHB_HUMAN | P07195 | LDHB | L-lactate dehydrogenase B chain |
| 212 | CDC42_HUMAN | P60953 | CDC42 | Cell division control protein 42 homolog |
| 213 | COF1_HUMAN | P23528 | COF1 | Cofilin-1 |
| 226 | LG3BP_HUMAN | Q08380 | LG3BP | Galectin-3-binding protein |
| 232 | ACLY_HUMAN | P53396 | ACLY | ATP-citrate synthase |
| 246 | FAS_HUMAN | P49327 | FAS | Fatty acid synthase |
| 245 | CD9_HUMAN | P21926 | CD9 | CD9 antigen |
| 268 | TCPD_HUMAN | P50991 | TCPD | T-complex protein 1 subunit delta |

**Table S2.** Funrich analysis of wound healing-associated proteins found in sEV samples (n = 3).

| **Biological process/pathway** | **% proteins** | **p-value** | **UniProt Acession Code** |
| --- | --- | --- | --- |
| neutrophil degranulation | 32,26 | 1,61E-09 | P05164; P00738; P27105; Q02413; P49913; P15924; P68871; P14923; P05109; P46976; |
| negative regulation of apoptotic process | 16,13 | 0,00056 | P02768; P05164; P62987; P08758; P09525; |
| defense response to bacterium | 16,13 | 3,61E-06 | P05164; P00738; P49913; P05109; P81605; |
| receptor-mediated endocytosis | 16,13 | 4,35E-06 | P02768; P00738; P69905; P02790; P68871; |
| antimicrobial humoral response | 12,90 | 1,17E-06 | P59666; P49913; P05109; P81605; |
| defense response to fungus | 12,90 | 6,63E-08 | P05164; P59666; P05109; P81605; |
| innate immune response | 12,90 | 0,004317 | P62987; P59666; P49913; P05109; |
| keratinization | 9,68 | 0,002687 | Q02413; P15924; P14923; |
| hydrogen peroxide catabolic process | 9,68 | 3,12E-06 | P05164; P69905; P68871; |

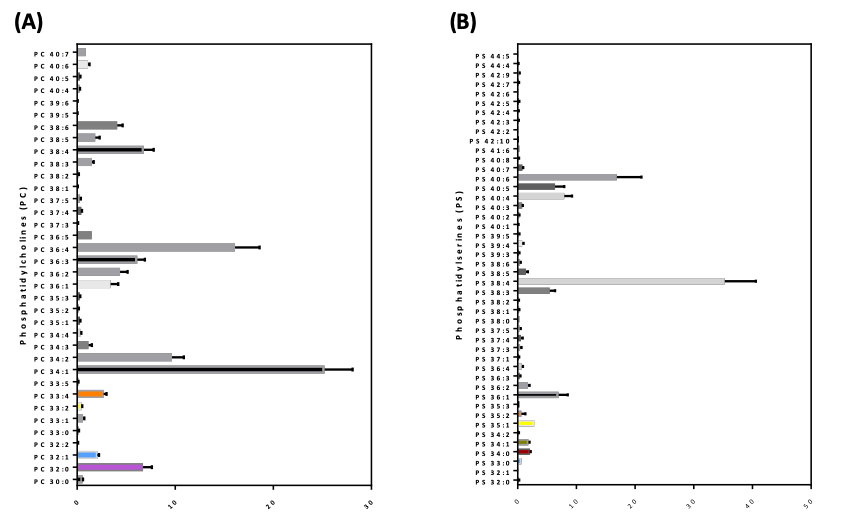

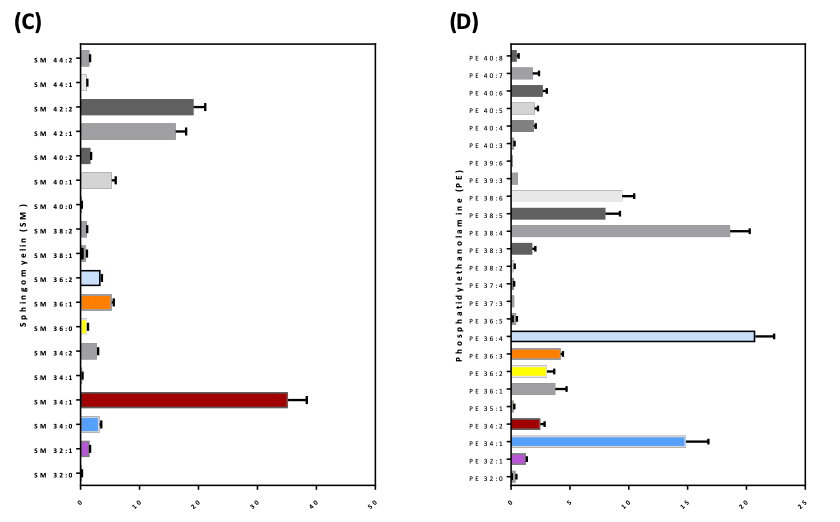

**Figure S4: Lipid content of sEV samples.** Quality of **(A)** phosphatidylcholines (PC), **(B)** phosphatidylserines (PS), **(C)** sphingomyelin (SM), and **(D)** phosphatidylethanolamines (PS) present in sEV samples (n = 5). Values are mean ± SEM.

**Table S3.** Characteristics of sEV isolated from umbilical cord blood mononuclear cells.

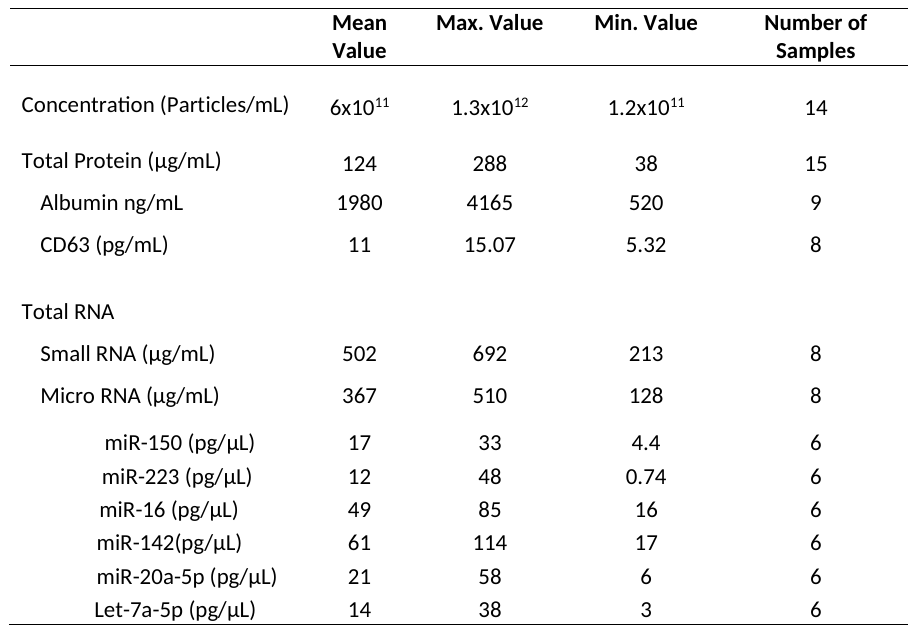

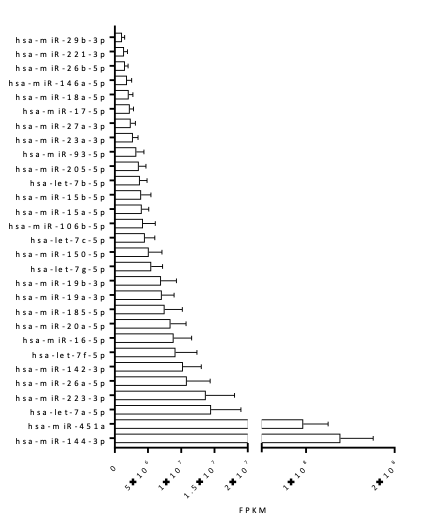

**Figure S5: Characterization of microRNAs in sEV.** Common miRNA identified in sEV samples (n = 6), organized by isoform-level relative abundance in fragments per kilobase of exon model per million mapped fragments (FPKM). Values are mean ± SEM.

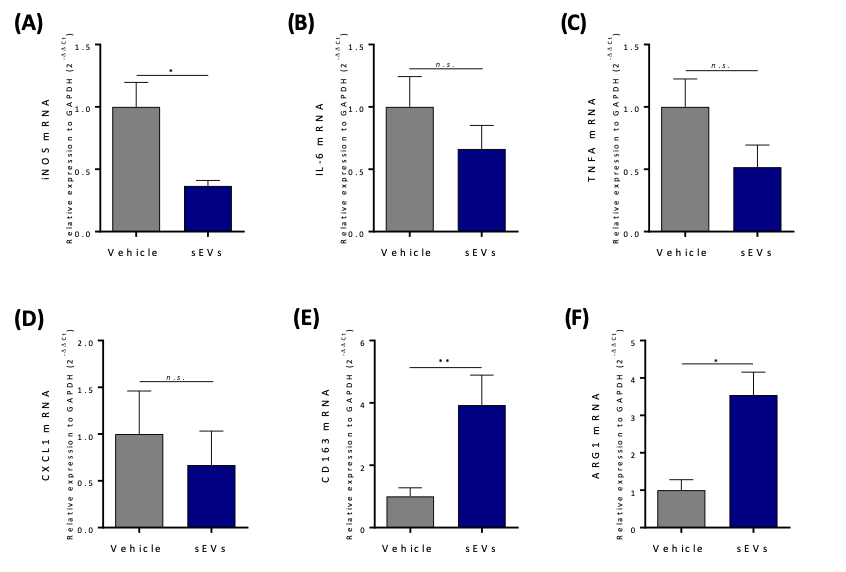

**Figure S6: Gene expression of chronic wounds after sEV treatment.** Fold change of genes associated with inflammatory response or macrophage phenotype. **(A)** Inducible nitric oxide synthase (iNOS), **(B)** IL-6, **(C)** TNF-α, **(D)** CXCL1, **(E)** CD163, and **(F)** arginase-1 (n = 3). All values are mean ± SEM. *n.s.* = non-significative, *p<0.05, **p<0.01.

**Table S4.** Funrich analysis of differentially regulated genes in wounds treated with sEV versus PBS, at days 3 and 15 (n = 3, p < 0.05).

| **Biological Process** | **Day 3** | | **Day 15** | |
| --- | --- | --- | --- | --- |
|  | **# genes** | **% of total** | **# genes** | **% of total** |
| regulation of transcription | 176 | 13.96 | 36 | 6.33 |
| angiogenesis | 19 | 1.51 | 17 | 2.99 |
| apoptotic process | 104 | 8.25 | 24 | 4.22 |
| cell adhesion | 31 | 2.46 | 29 | 5.10 |
| cell differentiation | 60 | 4.76 | 23 | 4.04 |
| cell proliferation | 87 | 6.90 | 47 | 8.26 |
| ECM organization | 3 | 0.24 | 11 | 1.93 |
| immune system process | 99 | 7.85 | 13 | 2.28 |
| inflammatory response | 95 | 7.53 | 8 | 1.41 |
| ion transport | 34 | 2.70 | 6 | 1.05 |
| metabolic process | 74 | 5.87 | 5 | 0.88 |
| oxidation-reduction process | 94 | 7.45 | 28 | 4.92 |
| translation | 45 | 3.57 | 18 | 3.16 |
| Wnt receptor signaling pathway | 16 | 1.27 | 14 | 2.46 |
| Total differentially expressed genes | 1261 | | 569 | |

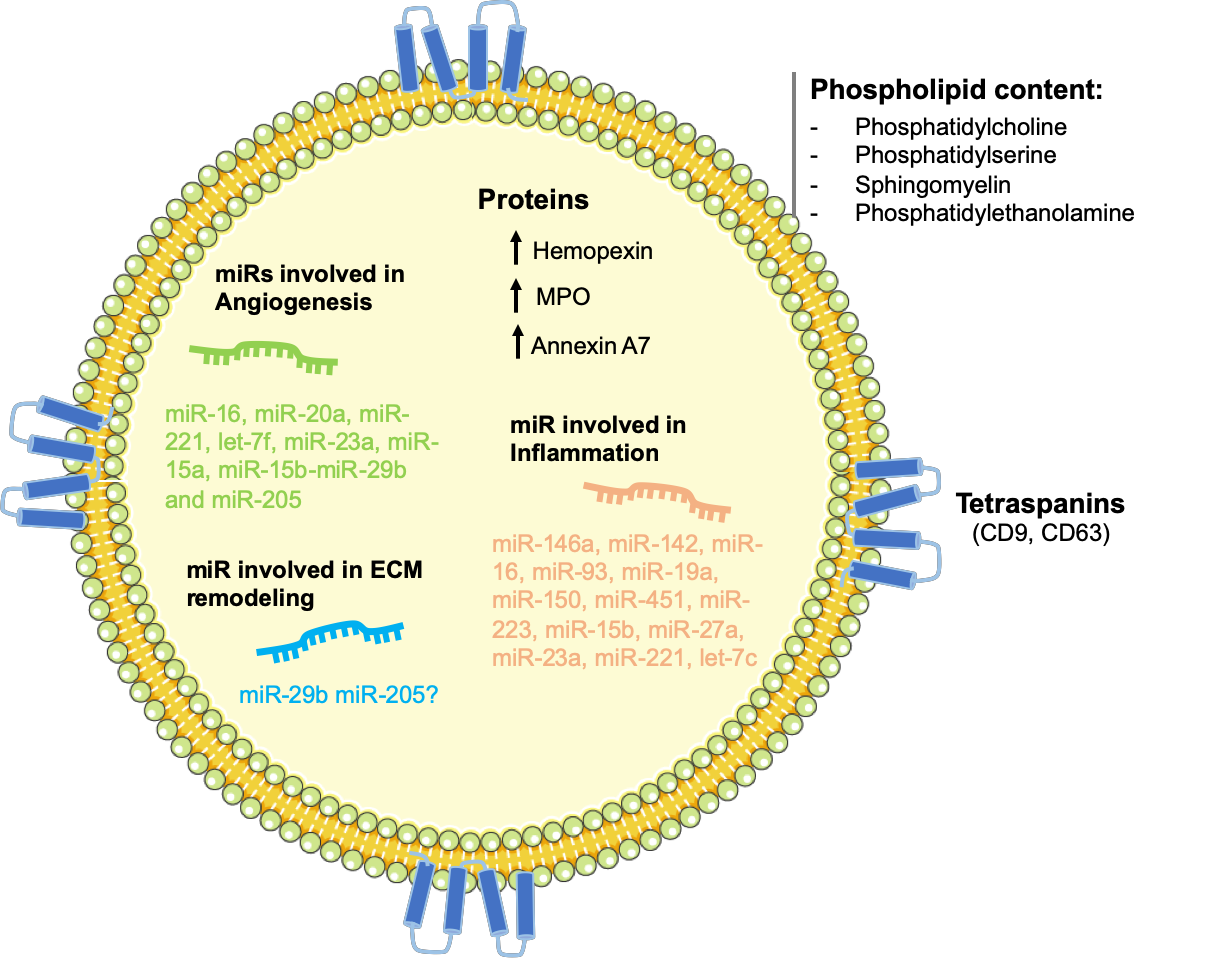

**Figure S7: Schematic illustration reflecting the composition of UCB-MNC-sEV and possible links to the mode of action in wound healing.**

Small Extracellular Vesicles (sEV) produced by Umbilical Cord Blood Mononuclear Cells (UCB-MNC) are rich sources of specific mRNAs, proteins and lipids, which are likely involved in regenerative processes, such as wound healing. Here, were isolated UCB-MNC-sEV with an optimized methodology, which increases yield and standardization, reduces production time and can easily be upscaled and GMP-compliant. This work evidences the therapeutic potential of sEV and focuses on optimizing production, bringing these vesicles closer to the clinics.

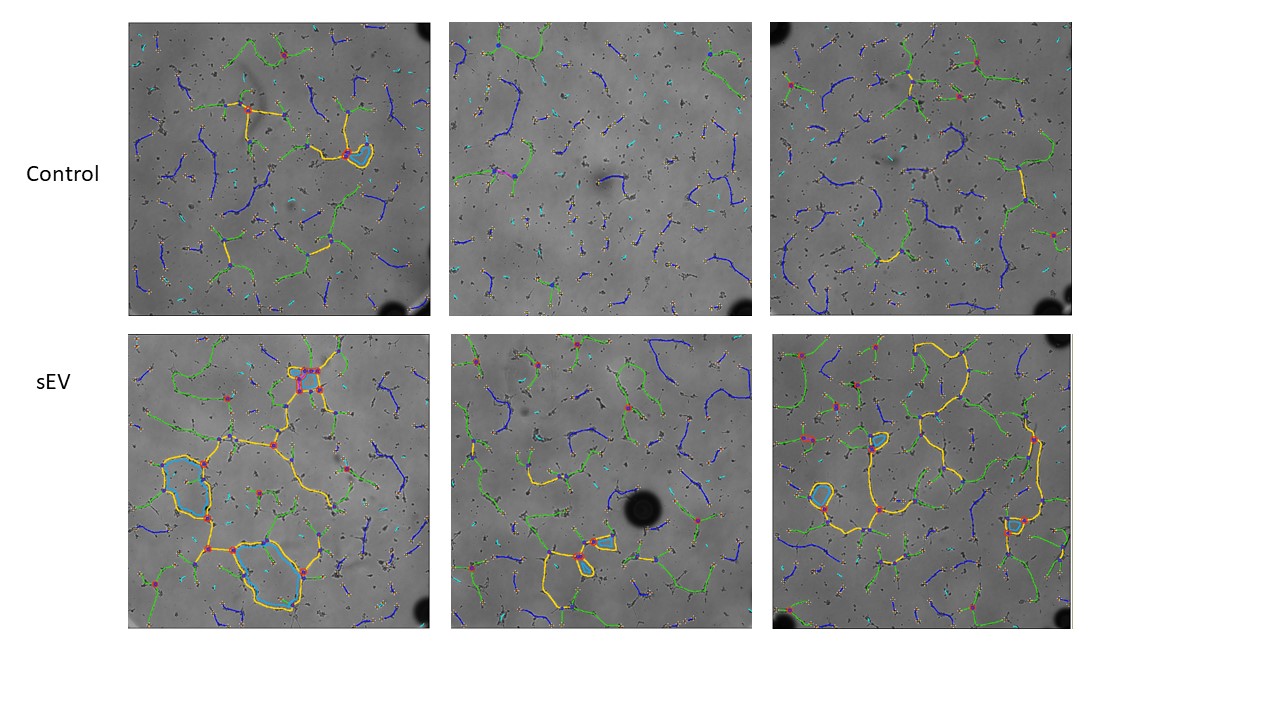

**Figure S8: Three representative images of Matrigel tube formation assay at 4h after treatment administration.** 10,000 endothelial cells/well were seeded on polymerized Matrigel (BD Biosciences, San Jose, CA) in a 96-well plate in complete endothelial media (ATCC, Manassas, VA). After 24h, the media was replaced with a starvation media, either alone or supplemented with 1x10^10^ sEV/mL. Cells were photographed, after 4 hours using an InCell microscope (GE Healthcare, Chicago, IL). In sEV treated samples, a higher endothelial cell reorganization to form tube or capillary-like structures is visible. After image processing, increased number of nodes and meshes was identified as well as higher total tube length. Image analysis software marked linear and branched segments as blue and green, respectively. Yellow lines represent master segments and light blue represent meshes. Nodes are represented in red.

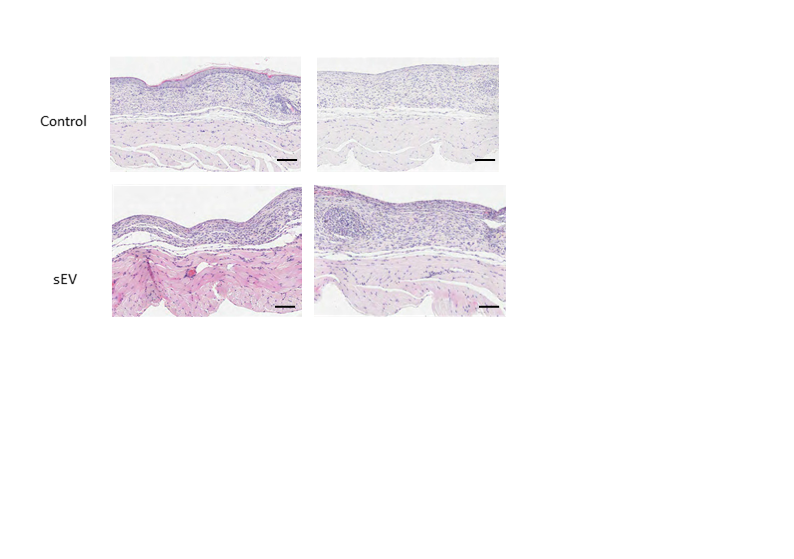

**Figure S9: Two representative images of mice skin biopsies.** On post-wounding day 20, tissues were harvested, fixed and embedded in paraffin wax. Representative sections (from the centre of each wound) were stained with Haematoxylin & Eosin - to facilitate measurement of granulation tissue depth, wound contraction and wound healing progression. Histologic analysis was blindly performed by a CICA biomedical pathologist. Scale bar: 100µm
